# Common variant burden contributes significantly to the familial aggregation of migraine in 1,589 families

**DOI:** 10.1101/226985

**Authors:** P. Gormley, M.I. Kurki, M.E. Hiekkala, K. Veerapen, P. Häppölä, A. Mitchell, D. Lal, P. Palta, I. Surakka, M.A. Kaunisto, E. Hämäläinen, S. Vepsäläinen, H. Havanka, H. Harno, M. Ilmavirta, M. Nissilä, E. Säkö, M-L. Sumelahti, J. Liukkonen, M. Sillanpää, L. Metsähonkala, P. Jousilahti, V. Anttila, V Salomaa, V. Artto, M. Färkkilä, 23andMe Research Team, The International Headache Genetics Consortium (IHGC), H. Runz, M.J. Daly, B.M. Neale, S. Ripatti, M. Kallela, M. Wessman, A. Palotie

**Author notes:** These authors contributed equally (P. Gormley, M.I. Kurki, M.E. Hiekkala). Corresponding author: Aarno Palotie.

## Abstract

It has long been observed that complex traits, including migraine, often aggregate in families, but the underlying genetic architecture behind this is not well understood. Two competing hypotheses exist, emphasizing either rare or common genetic variation. More specifically, familial aggregation could be predominantly explained by rare, penetrant variants that segregate according to Mendelian inheritance or rather by the sufficient polygenic accumulation of many common variants, each with an individually small effect. Some combination of both common and rare variation could also contribute towards a spectrum of disease risk.

We investigated this in a collection of 8,319 individuals across 1,589 migraine families from Finland. Family members were individually diagnosed by a migraine-specific questionnaire with either migraine without aura (MO, ICHD-3 code 1.1, n=2,357), migraine with typical aura (ICHD- 3 code 1.2.1, n=2,420), hemiplegic migraine (HM, ICHD-3 code 1.2.3, n=540), or no migraine (n=3,002). For comparison, we used population-based migraine cases (n=1,101) and controls (n=13,369) from the FINRISK study. The disease status of FINRISK individuals was assigned based on health registry data from outpatient clinics and/or prescription medication. All individuals were genotyped on the Illumina^®^ CoreExome or PsychArray chip platforms and imputed to a Finnish reference panel of 6,962 haplotypes. Polygenic risk scores (PRS), representing the common variant burden in each individual, were calculated using weights from the most recent large-scale genome-wide association study of migraine. To account for family structure in our analyses, we used a mixed-model approach, adjusting for the genetic relationship matrix as a random effect.

We found a significantly higher common variant burden in familial cases of migraine (for all subtypes, measured by the odds ratio [OR] per standard deviation [SD] increase in PRS; OR = 1.76, 95% CI = 1.71-1.81, P = 1.7×10^−109^) compared to cases from a population cohort (OR = 1.32, 95% CI = 1.25-1.38, P = 7.2×10^−17^) when using the population controls as a reference group. The highest enrichment was observed for HM (OR = 1.96, 95% CI = 1.86-2.07, *P* = 8.7×10^−36^) and migraine with typical aura (OR = 1.85, 95% CI = 1.79-1.91, *P* = 1.4×10^−86^) but enrichment was also present for MO (OR = 1.57, 95% CI = 1.51-1.63, *P* = 1.1×10^−48^). Comparing within cases, there was no significant difference in common variant burden between the migraine with aura subtypes, HM and migraine with typical aura (OR = 1.09, 95% CI = 0.99-1.19, *P* = 0.09), but both showed significantly higher enrichment compared to MO (OR = 1.28, 95% CI = 1.17-1.38, *P* = 7.3×10^−7^, and OR = 1.17, 95% CI = 1.11-1.23, *P* = 4.62×10^−5^, respectively). Additionally, we found that higher common variant burden corresponded to earlier age of headache onset (OR per SD increase in PRS for 3,631 cases with onset before 20 years old compared to 1,686 cases with onset later than 20 years old; OR = 1.11, 95% CI = 1.05-1.18, *P =* 8.3×10^−4^). FINRISK population cases identified from national health registry data were found to have lower common variant burden in comparison to the familial migraine cases (OR = 1.32, 95% CI = 1.25-1.38, *P =* 6.8×10^−17^), unless the individuals had attended both a specialist clinic and also received prophylactic migraine treatment (OR = 1.70, 95% CI = 1.53-1.88, *P =* 3.9×10^−9^). Finally, although rare variants have been suggested as the primary cause for familial hemiplegic migraine (FHM), we found only four out of 45 sequenced FHM families (8.9%) with a pathogenic mutation in one of the known risk genes.

In summary, our results demonstrate a substantial contribution of common polygenic variation to familial aggregation in migraine, comparable to both controls and that observed in migraine cases from a population cohort. The findings also suggest that individuals with migraine aura symptoms (either typical aura, which is mostly visual, or rare motor aura) tend to have higher common variant burden on average supporting the polygenic model also in these migraine subtypes.

## Introduction

Familial aggregation in chronic diseases is well-known but its background is not well understood ^1^. One hypothesis has been based on the Mendelian viewpoint that segregating, highly penetrant variants would strongly contribute to the familial nature of the disease. Linkage studies have had modest success in identifying highly penetrant disease variants; on the other hand, the numerous established genetic loci from genome-wide association studies (GWAS) rarely co-reside within linkage peaks. So far, most of the whole-exome sequencing (WES) and whole-genome sequencing (WGS) studies have been underpowered to shed light on the question of why common diseases aggregate in families. Studies in familial dyslipidemias have shed some light on this, by demonstrating that both rare (penetrant) and common (less penetrant) variants have been associated to specific lipid traits ^2,3^.

Migraine is an example of a common disease that can aggregate in families ^4^. It is one of the most common brain disorders worldwide, affecting approximately 15-20% of the adult population in developed countries ^5^. Therefore migraine studies may facilitate collection of the large sample sizes required to reveal some of the mechanisms of familial aggregation.

One third of migraine patients experience additional neurological symptoms during attacks, called aura (migraine with aura, MA, ICHD-3 code: 1.2). These can occur in rare forms called hemiplegic migraine (HM, ICHD-3 code: 1.2.3), typified by severe symptoms of motor aura that can be either familial (FHM, ICHD-3 code: 1.2.3.1) or sporadic (SHM, ICHD-3 code: 1.2.3.2). Alternatively, the more common form is usually accompanied by a visual aura, called migraine with typical aura (ICHD-3 code: 1.2.1). Migraine that occurs without any aura symptoms is the most common subtype and is called migraine without aura (MO, ICHD-3 code: 1.1). Each subtype is diagnosed according to the third edition of the International Classification of Headache Disorders (ICHD-3) criteria ^6^.

A Mendelian inheritance model in familial migraine has been supported by mutations in three ion-transporter genes *(CACNA1A* ^7^, *ATP1A2* ^8^, and *SCN1A* ^9^) identified by linkage studies and positional cloning of FHM families. However, mutations in these genes explain only a fraction of FHM/SHM cases ^10,11^ and none of the more common forms of migraine. Even in FHM these mutations vary in their penetrance, therefore, it is unlikely that penetrant mutations would entirely explain the observation that migraine is enriched in some families. Linkage studies in common forms of migraine have suggested several loci but no specific genes have been identified ^12^.

The polygenic nature of migraine is well documented by GWAS that have identified over 40 loci associated to common forms of migraine ^13-17^. Basic understanding of differences in the pathophysiology of these common forms (MA and MO) is limited. GWAS have identified many more common variant loci in MO than in MA ^13-17^, likely due to the larger sample sizes collected for MO, but clear differences in prevalence (MA = 5%, MO = 12%) could instead point towards differences in the genetic architecture and heterogeneity of these diseases.

We hypothesize that in addition to some rare, highly penetrant variants, accumulation of common variants with small individual effect sizes contribute to the familial forms of migraine. To study this, we constructed a polygenic risk score (PRS) from the most recent migraine GWAS consisting of approximately 59,000 cases and 316,000 controls after excluding all Finnish samples ^16^. We then investigated the contribution of common polygenic and rare variation to migraine in our large migraine family cohort consisting of 1,589 families from Finland totaling 8,319 individuals, including 540 HM, 2,420 migraine with typical aura, 2,357 MO, and 3,002 family members with no migraine. We observed an overall increased PRS in familial migraine cases compared to population-based cases and controls and clear differences of the common variant load across different migraine subtypes.

## Methods

### Sample collections

#### The migraine family collection

The families were collected over a period of 25 years from six headache clinics in Finland (Helsinki, Turku, Jyväskylä, Tampere, Kemi, and Kuopio) and through advertisements on the national migraine patient organization web page (www.migreeni.org). Geographically, family members are represented from across the entire country. The current collection consists of 1,589 families which included a complete range of pedigree sizes from small to large (e.g. 1,023 families had 1-4 related individuals and 566 families had 5+ related individuals, see **Supplementary Table S1** and **Figure S1**). It should be noted here that 455 individuals in the sample were single probands (i.e. unrelated cases without available affected relatives for analysis) but since they were ascertained in the same way as the other migraine families we have included them. Currently, the collection consists of 8,319 family members, of whom 5,317 have a migraine diagnosis based on the third edition of the established International Classification for Headache Disorders (ICHD-3) criteria ^6^. In about 50% of these affected individuals, the migraine attack is preceded by an aura phase. Another 3,002 family members were classified as having no migraines, including 1,557 individuals with no headache, 427 individuals with headache, 755 individuals with probable migraine, and 263 individuals with unknown diagnosis.

Migraine phenotype data was collected with a combination of individual interviews and an extensively validated Finnish Migraine Specific Questionnaire for Family Studies (FMSQFS ^18^). All participants were also asked to donate a blood sample. Over 200 variables were recorded, including information on the ICHD-3 symptoms, typical attack features, age of onset, other diseases, place of birth, etc. For the index patient in each family a neurologist performed a physical examination and sometimes other family members were examined as well. In all cases where the diagnosis was not clear from the questionnaire, a neurologist specialized in headache disorders interviewed the study subject. A summary of the sample characteristics of this family collection is shown in **Table 1 and Supplementary Table S2**.

**Table 1.** Sample characteristics of the FINRISK population-based sample (n = 14,470) and the Finnish migraine family collection (n = 8,319 individuals from 1,589 families). ICHD-3 codes are the classification codes from the 3^rd^ edition of the International Classification of Headache Disorders for each migraine subtype (note these classification criteria were not available for the FINRISK population sample). Mean PRS is the mean polygenic risk score calculated in each group.

| Sample group | ICHD-3 code | n | Mean age (SD) | Sex = Male (%) | Index case (%) | Mean PRS (SD) |
| --- | --- | --- | --- | --- | --- | --- |
| FINRISK cohort (all) | - | 14,470 | 46.44 (12.91) | 6,356 (43.9) | - | 0.02 (1.00) |
| • Controls | - | 13,369 | 46.96 (12.91) | 6,155 (46.0) | - | 0.00 (1.00) |
| • Any migraine | - | 1,101 | 40.09 (11.13) | 201 (18.3) | - | 0.26 (0.98) |
| □ triptans only | - | 812 | 39.65 (10.77) | 143 (17.6) | - | 0.22 (0.98) |
| □ outpatient only | - | 125 | 45.04 (13.26) | 37 (29.6) | - | 0.26 (0.98) |
| □ outpatient + triptans | - | 164 | 38.47 (10.11) | 21 (12.8) | - | 0.45 (0.93) |
| Family collection (all) | - | 8,319 | 44.56 (18.24) | 2,982 (35.8) | 1,568 (18.8) | 0.28 (1.02) |
| • No migraine | - | 3,002 | 44.67 (18.33) | 1,687 (56.2) | 85 ( 2.8) | 0.06 (1.01) |
| • Any migraine | 1 | 5,317 | 44.49 (18.19) | 1,295 (24.4) | 1,483 (27.9) | 0.41 (1.01) |
| □ migraine without aura (MO) | 1.1 | 2,357 | 44.85 (17.95) | 600 (25.5) | 561 (23.8) | 0.32 (0.99) |
| □ migraine with aura (MA) | 1.2 | 2,960 | 44.21 (18.38) | 695 (23.5) | 922 (31.1) | 0.48 (1.03) |
| □ migraine with typical aura | 1.2.1 | 2,420 | 44.35 (18.52) | 612 (25.3) | 636 (26.3) | 0.47 (1.03) |
| - typical aura with headache | 1.2.1.1 | 2,350 | 44.32 (18.50) | 575 (24.5) | 633 (26.9) | 0.47 (1.03) |
| - typical aura without headache | 1.2.1.2 | 70 | 45.09 (19.48) | 37 (52.9) | 3 ( 4.3) | -0.13 (0.94) |
| □ hemiplegic migraine (HM) | 1.2.3 | 540 | 43.58 (17.72) | 83 (15.4) | 286 (53.0) | 0.54 (0.99) |
| - familial hemiplegic migraine (FHM) | 1.2.3.1 | 197 | 44.83 (17.11) | 43 (21.8) | 53 (26.9) | 0.51 (1.00) |
| - sporadic hemiplegic migraine (SHM) | 1.2.3.2 | 343 | 42.86 (18.05) | 40 (11.7) | 233 (67.9) | 0.56 (0.98) |

#### FINRISK population-based cohort

FINRISK is a series of population-based health examination surveys carried out every five years since 1972 to monitor the risk of chronic diseases in Finland, as detailed elsewhere ^19^. Individuals in these cohorts have been prospectively followed for cardiovascular events and cause-specific death until 31^st^ December 2015 using annual record linkage with the Finnish National Hospital Discharge Register and the National Causes-of-Death Register. A total of 14,470 subjects from FINRISK were genotyped in five batches ^20^, having been randomly sampled from the full cohort, stratified by sex and cohort year (i.e. FINRISK 1992, 1997, 2002 or 2007 cohorts). A summary of the FINRISK sample characteristics is shown in **Table 1 and Supplementary Table S2**.

#### Finnish National Health Registry data for population-based cases

The population-based migraine cases within the FINRISK cohort were identified using Finnish National Health Registry data by two means: 1) From a specialist outpatient registry (from 1998 onwards) if an individual had received a migraine diagnosis (ICD-10 code: G43 or ICD-9 code: 346) either during a hospital visit (hospital discharge registry) or a speciality outpatient clinic visit (outpatient discharge registry) or 2) From a prescription drug purchase registry (from 1995 onwards) if an individual had been prescribed triptans at least twice (ATC codes under the N02CC category). This approach is likely to underreport migraine cases, particularly those with a sufficiently mild form of the disease so as to require neither triptan use nor visits to a hospital outpatient specialty clinic. Additionally, this data does not have symptom level information, so ICHD-3 based classification into migraine subtypes was not possible. Altogether 1,101 individuals fulfilling one of the above criteria were identified among a total of 14,470 study participants, giving a frequency of 7.6% of migraine cases.

### Genotyping and Quality Control

Genotyping was performed in seven batches on either the Illumina^®^ CoreExome or Illumina^®^ PsychArray, which share the Infinium^®^ HumanCore backbone including 480,000 variants in common. Samples from the migraine family collection were genotyped in two batches, one on the CoreExome and one on the PsychArray, with cases and controls distributed across both batches. Individual samples from the FINRISK cohort were genotyped in five batches, all on the CoreExome. A summary of these genotyping batches is provided in **Supplementary Table S3**. Before merging any batches we performed standard quality control procedures on each dataset individually, according to established GWAS protocols ^21^. Briefly, we excluded markers that exhibited high ‘missingness’ rates (>5%), low minor allele frequency (<1%), or failed a test of Hardy-Weinberg equilibrium (P<10^−6^). We also excluded individuals with high rates of heterozygosity (> 3 standard deviations from the mean), or a high proportion of missing genotypes (>5%). To control for any possible population stratification, we merged the genotypes from individuals passing QC with HapMap III data from European (CEU), Asian (CHB+JPT), and African (YRI) populations. We then performed a principal-components analysis on this combined data and excluded any population outliers not clustering with the other Finnish samples. We also performed a second principal-components analysis within each batch to ensure that cases and/or controls were clustering evenly together.

We then merged genotyping batches one-by-one and repeated the QC procedures described above on the merged dataset. To prevent any potential batch effects in the merged data, we also excluded any markers that failed a test of differential missingness (P<10^−5^) between the merged batches. Furthermore, during each round of merging, we performed a pseudo-association analysis (using a logistic mixed-model for batches with related individuals) between samples from each batch to identify markers where the minor allele frequency deviated significantly between batches (P<10^−5^). Markers with significant deviation were subsequently removed.

Finally, for the FINRISK samples we additionally used identity-by-descent (IBD) estimates to remove any closely related individuals (proportion IBD > 0.185), as the goal was to use them as a set of independent population controls. We further calculated kinship coefficients between all individuals using the software KING ^22^ in order to estimate genetic relatedness and to correct or remove individuals causing clear pedigree errors in the family sample.

### Reference panel for genotype imputation

To impute missing genotypes into the merged dataset (migraine families and FINRISK) we created a Finnish population-specific reference panel derived from sequencing data generated as part of the Sequencing Initiative Suomi (SISu) project ^23,24^. The reference panel combined low-coverage (mean depth ~4.6x) WGS data and high coverage WES data described further below.

#### Finnish low-coverage (~4.6x) WGS reference dataset

Sample- and variant-level quality control for the data was done at the Wellcome Trust Sanger Institute and only 1,940 high-quality unrelated individuals and polymorphic autosomal PASS SNP variants were included in the reference panel. Additionally, SNPs in low-complexity regions and those with Hardy-Weinberg equilibrium (HWE) P-value < 10^−5^ (n=99,191) were removed leaving a total of 13,625,209 markers after quality control. The data was then phased using SHAPEIT2 with default options and effective population size of 11,418.

#### Finnish WES reference dataset

From the raw WES data, variants were filtered according to the following criteria: 1) Multiallelic variants were removed, 2) Genotypes with QC < 20 were set to missing, 3) SNPs with call rate < 95% were removed, and 4) Monomorphic markers were removed. In addition, samples that were also in the WGS panel (n=7) were excluded together with individuals whose genotyping rate was < 95% (n=43). After filtering steps, the reference panel contained 1,540 individuals and 3,008,675 markers. The data was phased using SHAPEIT2 using default options and effective population size of 11,418.

Finally, the two reference panels (WGS and WES) were combined during imputation of the FINRISK and migraine family data using the software IMPUTE2 and its option to merge reference panels (i.e. ‘-merge_ref_panels’ option). We treated all available haplotypes from the two reference panels as informative (i.e. set total number of haplotypes as 6,962 with parameters: ‘-k_hap 3882 3080’).

### Imputation

Following genotyping QC, phased haplotypes were estimated for each individual using the program SHAPEIT2 ^25^ and its duoHMM method to improve accuracy by refining the estimation to haplotypes that are consistent with the pedigree structure. For phasing we chose an effective population size of 11,418, a window size of three, and 200 states for fitting the model. Missing genotypes were then imputed into these haplotypes using the program IMPUTE2 ^26^ and a manually created Finnish reference panel described above. We chose an effective population size of 20,000. While all samples (migraine family cases, controls, and FINRISK population) were imputed together, we split chromosomes into chunks of 3Mb with 500kb buffer to reduce the computation.

### Statistical methods

#### Calculation of polygenic risk scores (PRS)

To calculate the migraine PRS, we used the SNP effect sizes estimated for common variants from a previously published GWAS of migraine in 375,000 individuals ^16^. To ensure there were no overlapping samples from our family collection or FINRISK cohort, we excluded all samples of Finnish descent (i.e. four cohorts; the Finnish MA, Health 2000, NFBC, and Young Finns) from the original 22 cohort GWAS (**Supplementary Table S4**). We then recalculated the SNP effect size estimates for migraine from the remaining 18 studies that were of other European origin (i.e. 57,471 cases and 305,141 controls) using a fixed-effects meta-analysis. We then took the intersection of variants from the migraine GWAS dataset that overlapped with the imputed variants from our combined dataset of the migraine family collection and the FINRISK population. Next, we reduced the list of intersecting variants to an independent set by performing LD-clumping (r^2^ < 0.1 within 500kb from the most significant variant in each locus) using PLINK ^27^. Finally, we chose a subset of SNPs (n = 38,872) from the list of independent variants with P-values below a threshold of 0.1 in order to capture most of the variation influencing migraine risk while excluding the remainder of variants that do not show even a modest association. We then calculated the PRS for each individual as a sum of these alleles, weighted by the effect size estimates from the migraine GWAS results.

#### Association Analyses

To account for the high degree of relatedness within our family sample, we used logistic mixed-models to adjust for the genetic relatedness matrix (GRM) as a random effect. We calculated the GRM after filtering to a set of independent LD-pruned common SNPs (minor allele frequency > 5% and SNP missingness < 3%) using the program PLINK ^27^ (parameter options: ‘-- maf 0.05 --geno 0.03 --make-rel square gz’). In addition to adjusting for the GRM as a random effect, we also adjusted for sex, age, age^2^, and age^3^ as fixed effects. We then tested if the PRS was associated with migraine phenotypes using a Wald test of one degree of freedom. All mixed models and Wald tests were implemented in the statistical software R using the GMMAT package ^28^. We adjusted for multiple testing using Bonferroni correction.

#### Estimation of variance explained

To estimate the variance explained by the PRS, we fitted the same logistic mixed-model described above, adjusted for relatedness using GRM (random effect variable) and additionally adjusted for sex, age, age^2^, and age^3^ (fixed effect variables). We then compared the full model (including the PRS) with the null model (with PRS variable excluded) and estimated the variance explained using Nagelkerke’s pseudo-R^2^ (**Supplementary Table S5**).

#### Polygenic Transmission Disequilibrium Test (pTDT)

To assess polygenic burden of migraine risk alleles over-transmission from parents to affected offspring, we used the Polygenic Transmission Disequilibrium Test (pTDT) method ^29^. The method is robust to the relatedness structure as it uses full trios within a family sample and calculates an expected distribution of polygenic risk scores for offspring based on the average PRS of the parents. This expected distribution for the offspring is then used to test deviations from the null hypothesis in the observed mean parent PRS distribution. To calculate the expected distribution, we first separated the families into nuclear trios for any migraine (n = 1,486 trios) and then further performed subset analyses for MO (ICHD-3 code: 1.1, n = 737 trios), migraine with typical aura (ICHD-3 code: 1.2.1, n = 571 trios), and HM (ICHD-3 code: 1.2.3, n = 188 trios). We tested this hypothesis separately for offspring that were both cases and controls using a two-tailed, one-sample *t*-test.

### Identification of families/individuals with known pathogenic variants

To identify any families/individuals that carry a known pathogenic variant, which we define as a rare, penetrant mutation contributing to the disease (similar to guidelines provided elsewhere ^30^), we have screened 302 (101 FHM and 201 SHM) HM patients for these mutations in the three known genes for FHM *(CACNA1A, ATP1A2,* and *SCN1A).* To extract a list of possibly pathogenic variants in these genes, the selected 302 HM patients were either WES (293 HM cases from 243 families) ^31^ or Sanger-sequenced (9 FHM cases from one family) ^31,32^. WES was performed at the Wellcome Trust Sanger Institute (Hinxton, Cambridgeshire, UK). Reads were aligned to the human reference sequence (hg19) and genotypes called together using GATK’s best practices pipeline. We then filtered out variants that were common in gnomAD (MAF>1% in either all populations combined, or in the Finnish population alone MAF>0.1%), were predicted to be benign by SIFT or POLYPHEN2, or that did not segregate with cases in the family collection ^31^.

## Results

In this study, we assessed the contribution of migraine-associated common genetic variation to a range of migraine subtypes in a collection of 1,589 families from Finland (**Supplementary Table S1 and Supplementary Figure S1**). We calculated the PRS for all 8,319 individuals in these families and 14,470 individuals from the FINRISK population by combining each individual’s genotypes with SNP-association summary statistics from previously reported migraine GWAS results (**Methods** and **Supplementary Figure S2**). We then used the migraine PRS to assess the relative polygenic load contributing to both prevalent subtypes of migraine (migraine without aura and migraine with typical aura), to the rare subtype (HM), and to other available subtypes (**Table 1**). The mean polygenic burden observed for each migraine subtype, the family members without migraine, and the FINRISK population controls are shown in **Figure 1**.

**Figure 1.**
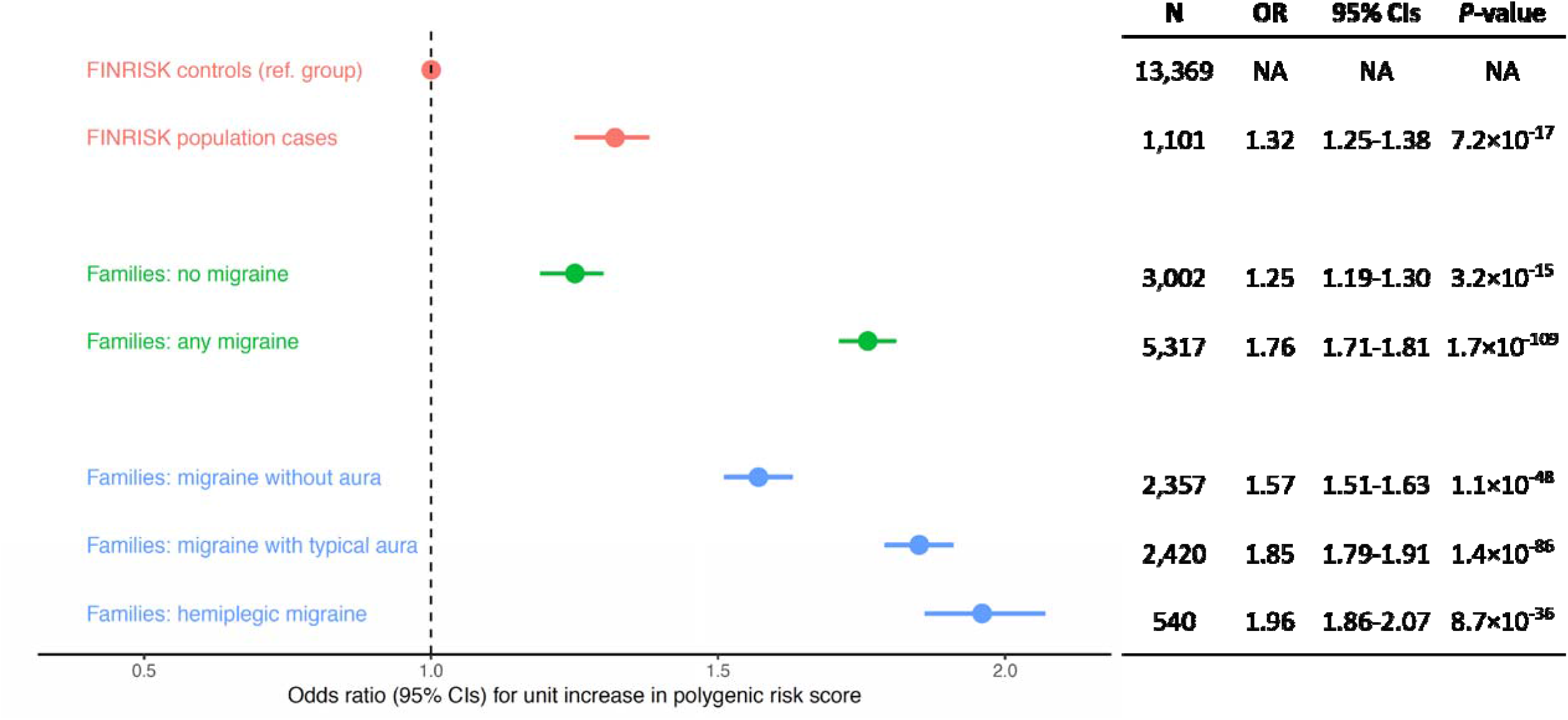
Odds ratio (OR) of a one standard deviation (SD) increase in Polygenic Risk Score (PRS) for different migraine phenotypes. The PRS was calculated using published migraine genome-wide association study (GWAS) data (N=375,000) from an independent set of 38,872 SNPs (with migraine GWAS association P-values < 0.1). ORs were calculated by comparing the PRS in each group to the 13,369 FINRISK population controls as a reference and using a logistic mixed-model adjusted for genetic relatedness, sex, and age. The 1,101 FINRISK population cases included any migraine cases identified from Finnish National Health Registry data. The groups marked “Families” are individuals from the Finnish Migraine Family Collection, including 3,002 with no migraine and 5,317 with any migraine. The 5,317 familial migraine cases were also broken down into specific subtypes, including; 2,357 migraine without aura (MO), 2,420 migraine with typical aura, and 540 hemiplegic migraine (HM).

### Common polygenic load is enriched across migraine subtypes

Compared to 13,369 FINRISK population-based controls, we found that the burden of common migraine-associated variation, measured via the PRS, was enriched across all of the migraine subtypes in the family collection, including rare forms of the disease. Using a logistic mixed model (adjusted for sex, age, and genetic relatedness) to test for association between the PRS as a continuous variable and each of the migraine subtypes we found that the PRS was associated with all migraine cases combined (OR = 1.76, 95% CI = 1.71-1.81, *P* = 1.7×10^−109^, **Table 1 and Figure 1**). We found the lowest enrichment for MO (OR = 1.57, 95% CI = 1.51-1.63, *P* = 1.1×10^−48^), compared to significantly higher enrichment for migraine with typical aura (OR = 1.85, 95% CI = 1.79-1.91, *P* = 1.4×10^−86^) and HM (OR = 1.96, 95% CI = 1.86-2.07, *P* = 8.7×10^−36^). From additional analyses using only cases to compare between migraine subtypes, there was no significant difference in common variant burden between the migraine with aura subtypes, HM and migraine with typical aura (OR = 1.09, 95% CI = 0.99-1.19, *P* = 0.09), but both showed significantly higher enrichment compared to MO (OR = 1.28, 95% CI = 1.17-1.38, *P* = 7.3×10^−7^, and OR = 1.17, 95% CI = 1.11-1.23, *P* = 4.62×10^−5^, respectively, **Supplementary Table S6**).

Investigating migraine with aura subtypes

We next looked at the four deeper-level subtypes of MA according to the ICHD-3 criteria. Two are subtypes of migraine with typical aura (ICHD-3 code: 1.2.1) called typical aura with headache (ICHD-3 code: 1.2.1.1) and typical aura without headache (ICHD-3 code: 1.2.1.2), and the other two are subtypes of HM, called FHM and SHM, where FHM is defined as those HM cases with at least one first- or second-degree relative that has also been diagnosed with HM (**Table 1**). We investigated if these sub-subtypes were any different in terms of their polygenic burden. We found that there was no significant difference between FHM and SHM (OR = 0.94, 95% CI = 0.72-1.16, *P* = 0.58, **Supplementary Table S6 and Supplementary Figure S3**). However, for migraine with typical aura we found that typical aura with headache cases had significantly higher polygenic burden than typical aura without headache (OR = 1.85, 95% CI = 1.45-2.36, *P* = 6.3×10^−7^, **Supplementary Table S6 and Supplementary Figure S3**). In fact, the PRS burden in cases of typical aura without headache was observed to be equivalently as low as the FINRISK population controls (OR = 0.85, 95% CI = 0.62 - 1.17, *P* = 0.20, **Supplementary Figure S3**).

### Increased risk between upper and lower PRS quartiles

To quantify the effect for individuals carrying the highest burden of risk alleles relative to the FINRISK population distribution, we separated individuals from the family collection into population-level quartiles of the PRS. We used the FINRISK population sample to calculate the cut-off values for individuals in the upper and lower quartiles of PRS and tested for enrichment between individuals in the highest and lowest quartiles of the distribution (**Supplementary Table S7**). Again, we observed the lowest enrichment in the MO subtype, where the mean PRS was estimated to be 2.2 times significantly higher than the mean PRS of the FINRISK population controls (OR = 2.2, 95% CI = 2.03-2.37, *P =* 1.4×10^−19^). The enrichment for migraine with typical aura was even higher, with mean PRS that was 3.0 times larger than the population mean (OR = 3.02, 95% CI = 2.85-3.20, *P =* 9.0×10^−35^). As before, in HM we observed the highest enrichment of common variation, with mean PRS that was 3.8 times significantly higher than the population mean (OR = 3.84, 95% CI = 3.52-4.15, *P =* 2.5×10^−17^).

### Comparing familial cases to population cases identified from the national health-registry

We observed that common variant burden measured via the PRS was higher in familial cases of any migraine subtype (n = 5,317) compared to population-based cases (n = 1,101) identified from FINRISK via health-registry data (OR = 1.26, 95% CI = 1.18-1.34, *P* = 3.2×10^−8^, **Supplementary Table S8**). Splitting this result by subtype, we found that the PRS in familial MO was modestly enriched compared to the population-based cases (OR = 1.13, 95% CI = 1.04-1.22, *P* = 0.0075), whereas migraine with typical aura and HM showed the highest PRS enrichment (OR = 1.31, 95% CI = 1.22-1.40, *P* = 4.6×10^−9^, and OR = 1.51, 95% CI = 1.37-1.64, *P* = 1.2×10^−9^ respectively).

We additionally found that migraine individuals within the family collection who had self-reported use of triptan medication had on average higher common variant burden compared with individuals who did not report use of triptans (OR = 1.12, 95% CI =1.06-1.19, *P* = 5.7×10^−4^), **Supplementary Table S9**). Among the FINRISK population cases, we did not observe any difference in PRS when comparing cases that were either triptan users (defined as purchasing triptans at least twice) or individuals who had visited a migraine outpatient clinic (OR = 1.06, 95% CI = 0.86-1.26, *P* = 0.56). However, individuals that had both visited a specialist outpatient clinic and used triptans had significantly higher PRS enrichment (OR = 1.70, 95% CI = 1.53-1.88, P = 3.9×10^−9^). In fact, this was similar enrichment to the PRS profile observed in the familial migraine cases (OR = 1.76, 95% CI = 1.71-1.81, P = 1.7×10^−109^). **Supplementary Figure S7**.

### Migraine-associated alleles are over-transmitted to affected offspring

As an additional method for investigating the contribution of polygenic load that is robust to relatedness in a family sample, we used the polygenic transmission disequilibrium test (pTDT, see **Methods**). Here we extracted nuclear trios from the family collection for four phenotypes (any migraine, MO, migraine with typical aura, and HM) and tested separately whether common migraine-associated alleles (as measured by the PRS) were disproportionately over-transmitted from parents to both affected and unaffected offspring. In support of the results from the mixed model, we found that affected offspring received a significantly higher transmission of common polygenic load from their parents than would be expected by chance alone (**Figure 2**). The over-transmission was observed in every migraine phenotype; any migraine, (P = 1.7×10^−13^, pTDT deviation = 0.16, 95% CI = 0.12 - 0.20), MO (P = 4.9×10^−4^, pTDT deviation = 0.10, 95%CI = 0.046 - 0.163), migraine with typical aura (P = 1.5×10^−6^, pTDT deviation = 0.17, 95%CI = 0.10 - 0.24), and HM (P = 3.7×10^−7^, pTDT deviation = 0.33, 95%CI = 0.21 - 0.45), see **Supplementary Table S10**. As expected, no over-transmission was observed for unaffected offspring (P>0.05). While the observed over-transmission of the PRS was higher for migraine with typical aura and HM compared to MO (consistent with the association results from the mixed-model above), the difference was not significant between migraine groups.

**Figure 2.**
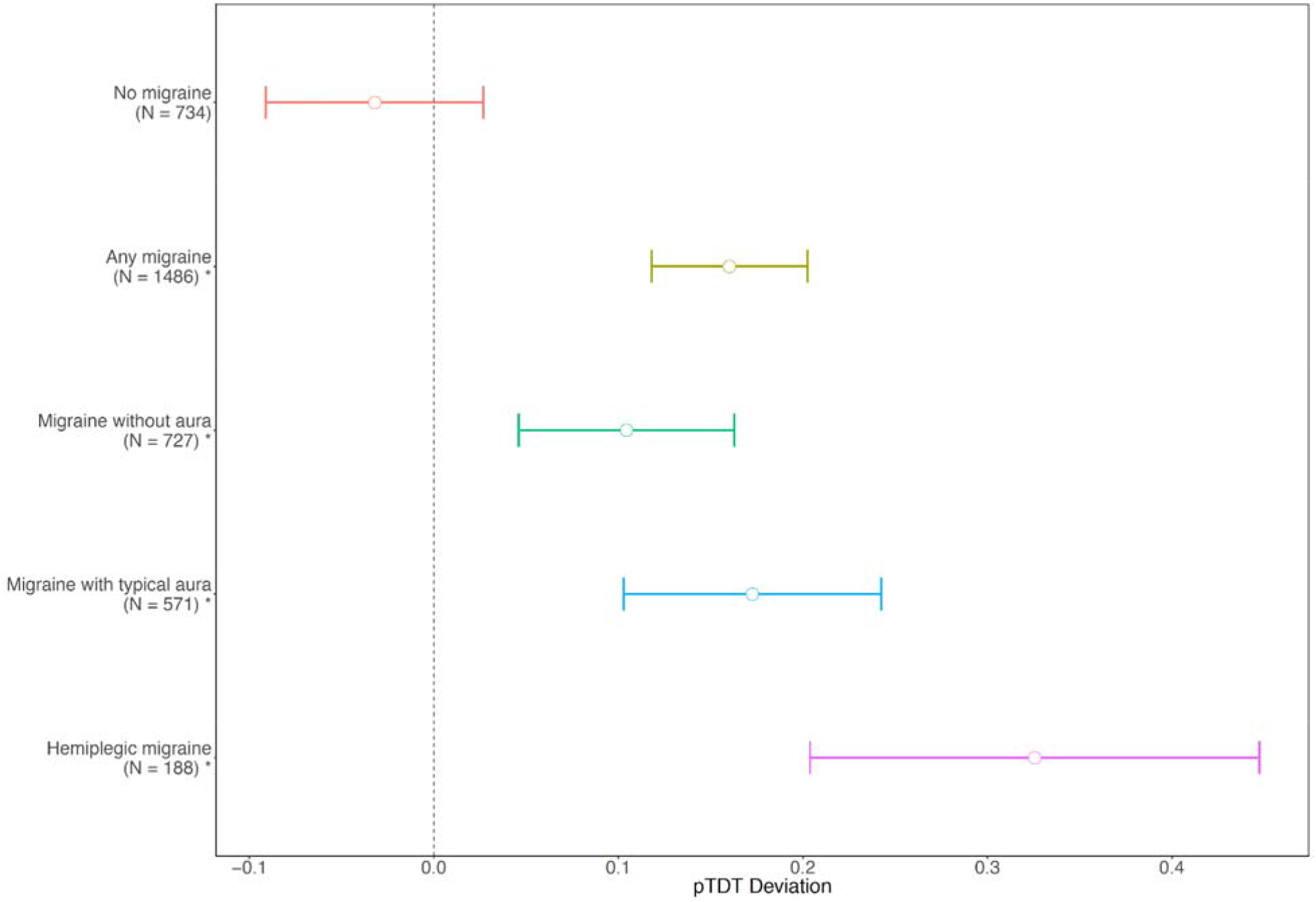
Polygenic Risk Score (PRS) transmission disequilibrium test (pTDT) in migraine subtypes. The Finnish migraine family collection was subset into whole trios and grouped based on disease status of the offspring. This included 734 trios where the offspring had no migraine, 1,486 trios where the offspring had any migraine. These trios were further divided into groups where the offspring had specific migraine subtypes, including 727 trios for migraine without aura (MO), 571 trios for migraine with typical aura, and 188 trios for hemiplegic migraine. The pTDT deviations were estimated based on deviations of the PRS observed in the offspring from the expected mean PRS to be transmitted by the parents. Groups that showed significant over-transmission are marked with ‘*’.

### Contribution of Mendelian SNPs and polygenic load to FHM

We examined the relative contribution of known pathogenic variants (see **Methods**) and polygenic load to FHM, using a subset of 74 families where familial aggregation of cases had been confirmed. From sequencing data on 101 FHM cases from 45 of these families, we have identified four families that carried a pathogenic, rare mutation in one of the three known FHM genes ^31,32^. Therefore out of the 45 sequenced families, 8.9% (4/45) could be potentially explained by a rare pathogenic mutation in one of the known FHM genes (**Table 2** and **Supplementary Table S11**). We have found no likely pathogenic mutations in these genes for any of the 201 (of 343) SHM individuals that were sequenced ^31^. We next investigated what proportion of cases from each migraine subtype were found in the extreme tails of the distribution of polygenic risk from the FINRISK population. The expected proportion of individuals in the upper quartile of risk is 25%, which was approximately what we observed for individuals with no migraine (26.7%), but found large deviations from expectation for the other phenotypes, including MO (36.2%), migraine with typical aura (41.4%), and HM (43.0%). Furthermore, of the 197 FHM cases from 74 families, 80 of these cases (40.6%) were in the highest quartile of polygenic risk (**Table 2** and **Supplementary Figure S4**).

**Table 2.**
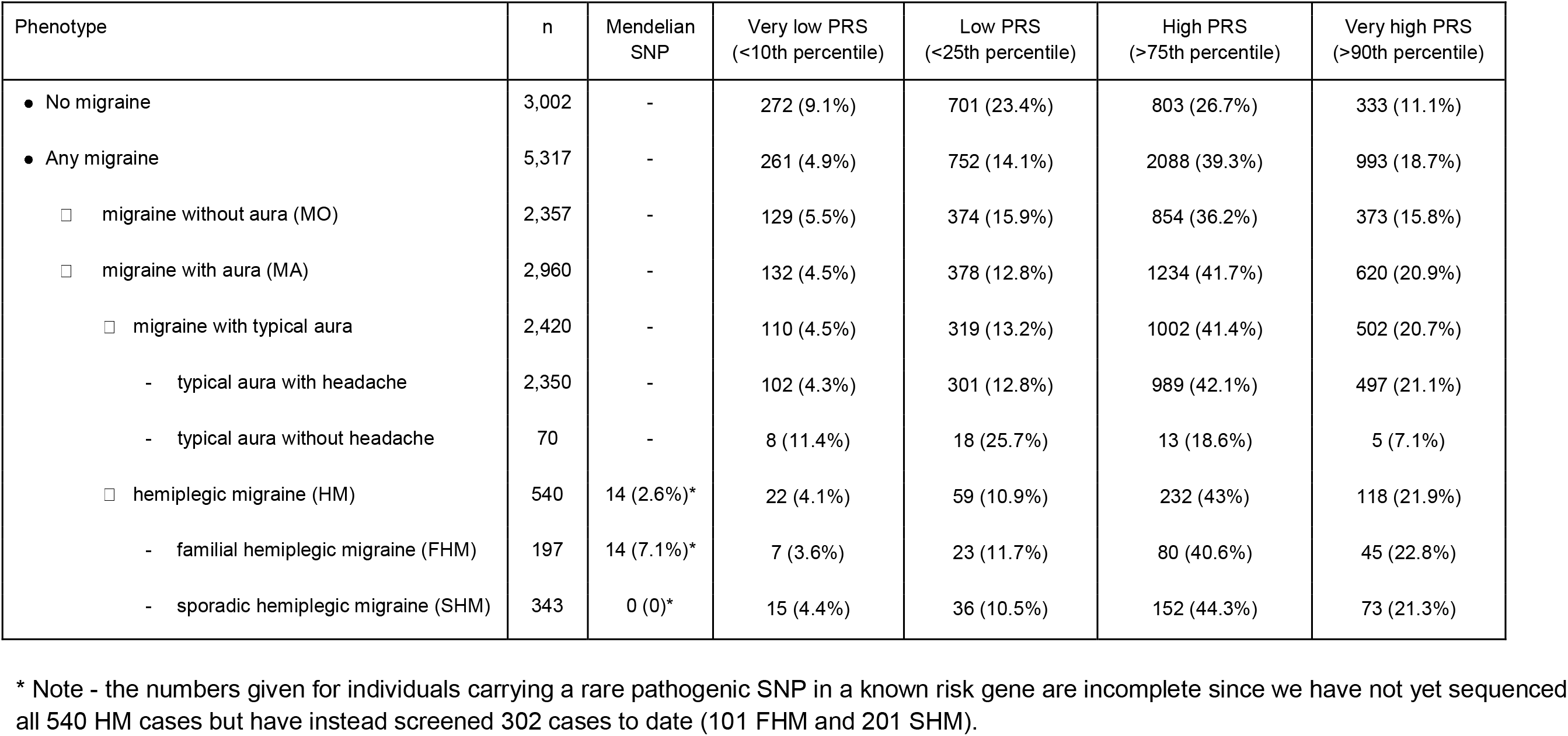
Number of affected individuals with a high Polygenic Risk Score (PRS) or a pathogenic Mendelian SNP. Affected individuals were assigned to high PRS category based on percentiles of the risk score distribution in the FINRISK population. 14 individuals from four families were categorized as having a previously reported pathogenic variant for FHM.

| Phenotype | n | Mendelian SNP | Very low PRS (<10th percentile) | Low PRS (<25th percentile) | High PRS (>75th percentile) | Very high PRS (>90th percentile) |
| --- | --- | --- | --- | --- | --- | --- |
| ● No migraine | 3,002 | - | 272 (9.1%) | 701 (23.4%) | 803 (26.7%) | 333 (11.1%) |
| ● Any migraine | 5,317 | - | 261 (4.9%) | 752 (14.1%) | 2088 (39.3%) | 993 (18.7%) |
| □ migraine without aura (MO) | 2,357 | - | 129 (5.5%) | 374 (15.9%) | 854 (36.2%) | 373 (15.8%) |
| □ migraine with aura (MA) | 2,960 | - | 132 (4.5%) | 378 (12.8%) | 1234 (41.7%) | 620 (20.9%) |
| □ migraine with typical aura | 2,420 | - | 110 (4.5%) | 319 (13.2%) | 1002 (41.4%) | 502 (20.7%) |
| - typical aura with headache | 2,350 | - | 102 (4.3%) | 301 (12.8%) | 989 (42.1%) | 497 (21.1%) |
| - typical aura without headache | 70 | - | 8 (11.4%) | 18 (25.7%) | 13 (18.6%) | 5 (7.1%) |
| □ hemiplegic migraine (HM) | 540 | 14 (2.6%)* | 22 (4.1%) | 59 (10.9%) | 232 (43%) | 118 (21.9%) |
| - familial hemiplegic migraine (FHM) | 197 | 14 (7.1%)* | 7 (3.6%) | 23 (11.7%) | 80 (40.6%) | 45 (22.8%) |
| - sporadic hemiplegic migraine (SHM) | 343 | 0 (0)* | 15 (4.4%) | 36 (10.5%) | 152 (44.3%) | 73 (21.3%) |

\* Note - the numbers given for individuals carrying a rare pathogenic SNP in a known risk gene are incomplete since we have not yet sequenced all 540 HM cases but have instead screened 302 cases to date (101 FHM and 201 SHM).

To assess the polygenic burden per family, we inspected the distribution of the median PRS from each of the 74 families with confirmed FHM cases (**Figure 3**). We found that over 44% of FHM families (33 out of 74) carried a common variant burden that was in the highest quartile of population risk. Additionally, only five of these 74 FHM families (6.8%) were found to be in the lowest quartile of population risk. Interestingly, the four apparently Mendelian families that carry a pathogenic rare mutation were not among the five families in the lowest quartile of risk and were instead spread across the distribution (in fact two families were in the highest PRS quartile, **Figure 3**), potentially indicating that other genetic and/or environmental factors also play a role in these families.

**Figure 3.**
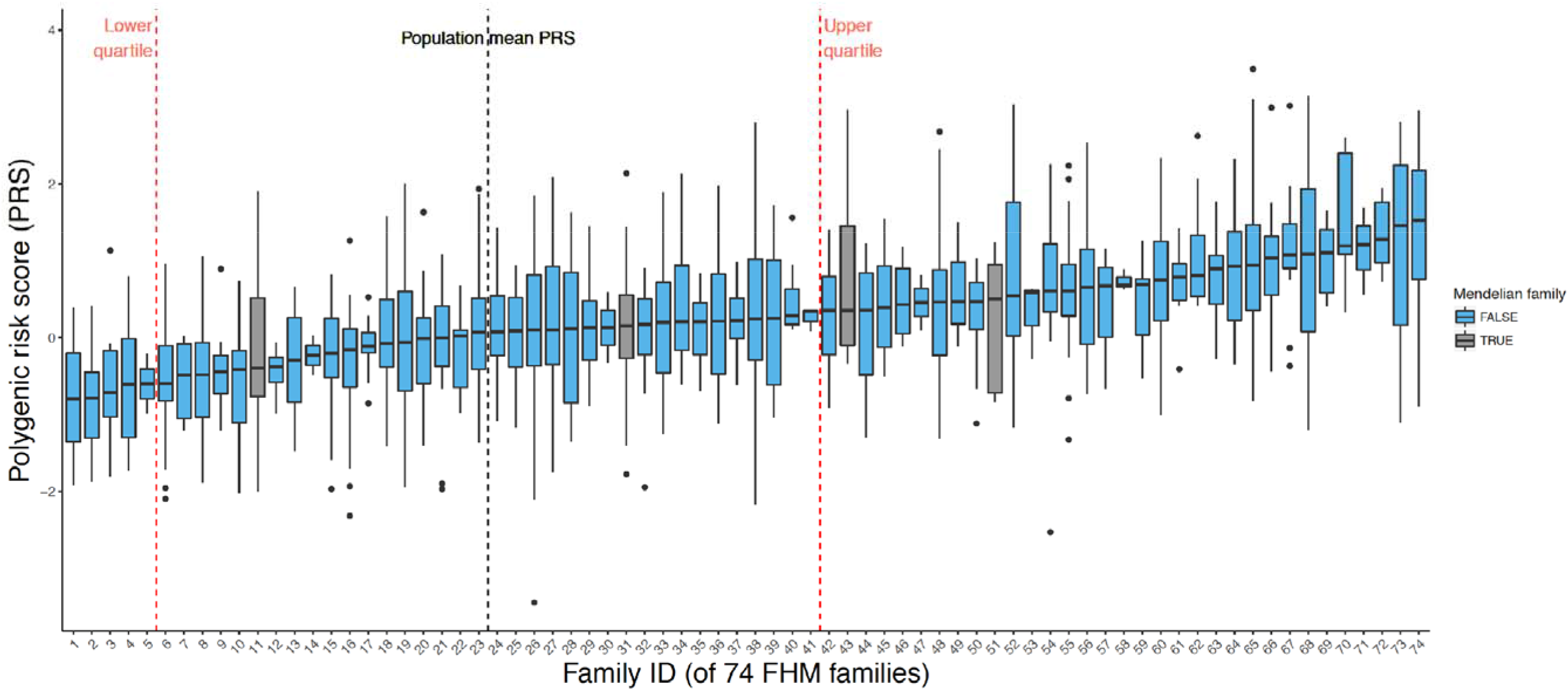
Boxplots per family of the Polygenic Risk Score (PRS) in each of the 74 FHM families. The horizontal axis is Family ID ordered by the mean PRS of each family. The four families carrying a known pathogenic/Mendelian SNP are highlighted in grey. The black vertical dashed line is the mean PRS of the FINRISK population controls. The red vertical dashed lines indicate the lower and upper quartiles of PRS of the FINRISK population controls.

### Relationship between age of onset and polygenic load

For a subset of individuals (n = 4,930) in the family collection we had information on age of onset of the migraine headache. We used this information to assess whether polygenic load was associated with age of onset. We grouped these 4,390 individuals with onset data into age of onset bins (0 to 10 years old [n = 1,295], over 10 to 15 years old [n = 1,402], over 15 to 20 years old [n = 990], and over 20 years old [n = 1,243]) and estimated whether the mean PRS in each bin was significantly different (**Figure 4**). Additionally, data from all cases (n=5,317) was available on whether onset of headaches occurred before or after 20 years old, so we compared the PRS between these two groups using a logistic mixed model. We found that the mean PRS was significantly higher in migraine cases where headache onset occurred before 20 years of age compared to individuals with later onset (OR = 1.11, 95% CI = 1.05-1.18, *P* = 8.2×10^−4^, **Supplementary Table S9**).

**Figure 4.**
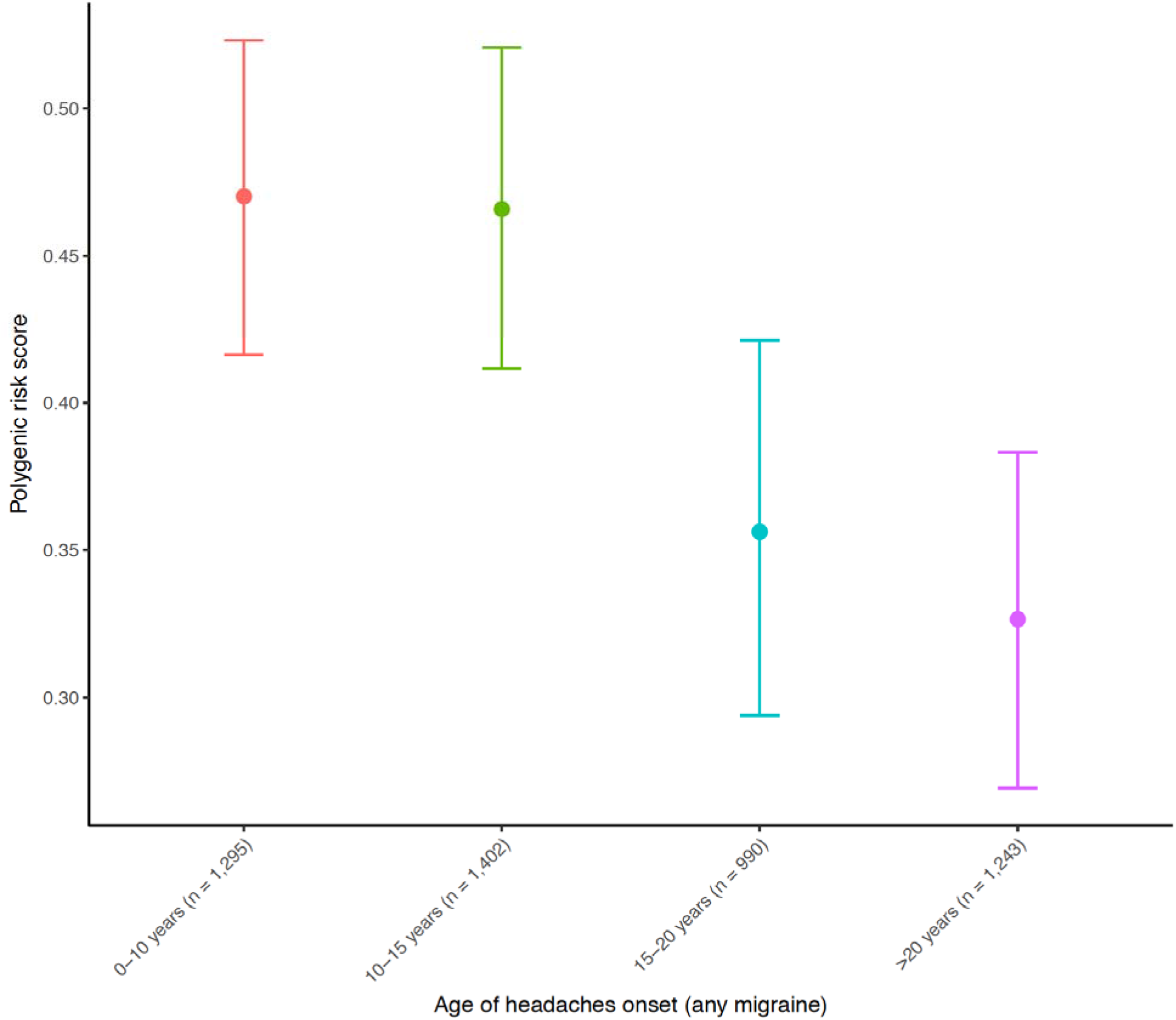
Mean Polygenic Risk Score (PRS) stratified by age of onset of headaches in migraine individuals. An increase in PRS corresponds to earlier age of onset of migraine headache. Means and 95% CIs were estimated within the family collection using bootstrap resampling (10,000 replicates) within each age of onset bin.

### Higher PRS is associated with higher rate of clinical diagnostic symptoms

We used nine diagnostic criteria for migraine (attack length > 4 hours, unilaterality, pulsation, moderate/severe intensity, aggravation by physical exercise, nausea, vomiting, phonophobia, and photophobia, **Supplementary Figure S5**) to test if the PRS was specifically associated with any of these individual criteria. As these data were obtained from some family members by questionnaire, only strict yes answers were interpreted as cases, whereas missing answers were interpreted as no-answers. We found that increased PRS predicted a higher rate of 8 out of 9 diagnostic symptoms (**Supplementary Table S12**), with severity of headache showing the largest effect size (OR = 1.29, 95% CI = 1.20-1.39, *P* = 1.1×10^−7^). Also strongly associated were ‘attack length greater than 4 hours’ (OR = 1.23, 95% CI = 1.16-1.29, *P* = 4.7×10^−9^) and photophobia (OR = 1.23, 95% CI = 1.14-1.32, *P* = 5.8×10^−6^). Notably, the only diagnostic criteria not associated with the polygenic risk score was headache pulsation (OR = 1.04, 95% CI = 0.98-1.10, *P* = 0.17).

## Discussion

Our results show that common polygenic variation, as measured via the PRS, significantly contributes to the familial aggregation of migraine. PRS enrichment in families was observed in both common and rare subtypes of familial migraine compared to both population controls and to population cases of migraine. There were relatively large differences in the PRS burden observed between different migraine sub-categories. The polygenic burden was higher for MA compared to those individuals that do not suffer any migraine aura symptoms. Individuals with migraine with typical aura were 3.0 times more likely to be in the top 25% of the population PRS compared to the bottom 25% of the distribution, while individuals with MO were only 2.2 times more likely. Strikingly, individuals with HM were observed to be enriched with the highest polygenic burden, where they were 3.8 times more likely to be in the upper 25% of population risk score. This is particularly interesting considering that FHM has previously been considered to be primarily driven by rare variants of large effect. Our results confirm that common variants identified by GWAS in populations play a considerable role in rare forms of migraine with aura (both FHM and SHM) and suggest that a large proportion of the disease risk in HM cases can be significantly explained by common polygenic variation, rather than solely by highly penetrant, rare variation.

In addition to showing that familial cases of migraine have higher polygenic burden on average compared to population controls, we also showed, using the pTDT approach, that offspring with migraine have inherited a higher burden of common polygenic variation associated with migraine than would be expected by chance alone. Together, these two methods produce results that are robust to genetic relatedness of individuals within the sample. Therefore, we have validated by two independent statistical methods (mixed-model and pTDT) our result that common polygenic variation associated with migraine significantly contributes to the familial aggregation of both prevalent and rare subtypes of migraine.

The National Health Register system provides data from every hospital- and out-patient visit and every prescription drug purchase of every citizen. This data was used to sub-categorize migraine cases in the population-based FINRISK sample. In addition to the formal ICD-code based migraine subtype definition, the registries enabled us to sub-categorize patients based on their use of the health care system. While a large fraction of migraine patients in the Finnish health care system are treated in primary care, the more complicated patients tend to be referred to secondary and tertiary treatment units, like hospital neurology outpatient clinics. Interestingly, migraine cases that had visited a specialist outpatient clinic and additionally had purchased triptans had the highest PRS, which was as high as the mean value of familial migraine cases ascertained from the clinics. This finding is consistent with recent observations in other traits, for example hyperlipidemias, where familial dyslipidemic cases were observed to have similarly high PRS compared to hyperlipidemia cases in the FINRISK population cohort ^3^. These findings suggest that severe cases identified from a population cohort, in terms of their polygenic profile, can be genetically similar to familial cases, and that familial aggregation mighi just be a reflection of a cumulative effect of many common variants.

Furthermore, we attempted to characterize the proportion of FHM cases that could be explained by rare pathogenic variants in the three known FHM genes. We identified only four out of 45 sequenced FHM families (8.9%) with cases that carried one of these variants, and 0 out of 201 sequenced SHM individuals ^31^. While it is possible that more pathogenic, rare variants for FHM are yet to be discovered, it is striking that so few known variants could be identified in our large family collection (n = 302 sequenced HM cases from 1,589 families). Together, with the observation of significant polygenic burden also seen in individuals with FHM and SHM (including over 40% of FHM cases from 74 families that were in the highest quartile of population polygenic risk), it is likely that a large proportion of the risk for these rare migraine phenotypes are explained by a higher burden of common polygenic variation, with HM falling on the high end of a spectrum of disease liability, possibly in some instances combined with rare variants of larger effect.

Another debate in the field where these results can potentially shed light on is the question of whether FHM and SHM are different diseases. Both diseases present with similar symptoms in the clinic, and out of the three known FHM genes, mutations in *CACNA1A* and *ATP1A2* have also been detected in SHM; however, the genetic factors influencing most cases remain unexplained ^33^. It has been speculated previously that due to the sporadic nature, SHM could be primarily driven by pathogenic *de novo* mutations, whereas FHM would instead be driven by pathogenic rare variants that segregate in families. However, our results show that both FHM and SHM cases combined carry the highest burden of common polygenic variation out of all migraine subtypes, and that there is no significant difference between the polygenic burden comparing between FHM and SHM. This suggests that these two migraine phenotypes share a similar genetic etiology and implicates the inheritance of a high common variant load by chance in sporadic forms of HM, and a high familial load of common variation in the aggregation of FHM in families. This is further supported by our observation that HM patients often have affected relatives with the common forms of migraine ^31^.

By contrast, for the more prevalent forms of MA, we found that there was a significant difference in common polygenic burden between the migraine with typical aura subgroups, typical aura with headache (ICHD-3 code: 1.2.1.1) and typical aura without headache (ICHD-3 code: 1.2.1.2). While the typical aura with headache group showed a similarly high polygenic burden compared with the more rare forms of MA (i.e. both FHM and SHM), and was not significantly different from these groups, we observed that the typical aura without headache group looked very different and carried a substantially lower common polygenic burden relative to the other MA subtypes. The contribution of the PRS (in terms of effect size) to this migraine phenotype was in fact no different than controls from the population (**Supplementary Figure S3**). As such, one might speculate that much of the common variation captured by the PRS is influencing genes specifically related to the etiology involved in the head pain characteristics of migraine, but more investigations are needed to determine this.

Interestingly, migraine cases in the family sample who experienced earlier age of onset of headaches tended to carry a higher polygenic burden of common migraine risk alleles on average. This association was observed for all migraine types and is consistent with similar findings in other complex disorders ^34^, as well as previous hypotheses that suggest that migraines that have earlier onset (before 15 - 20 years old) have a higher genetic burden compared to migraines that begin later in life, where some combination of genetic plus environmental factors (e.g. stress, diet, medications, general health, and other stimulants) may play a larger role.

Finally, when looking at nine diagnostic symptoms of migraine, we found that higher polygenic risk scores were associated with an increased risk for eight out of nine symptoms. The only symptom not associated with higher polygenic load was headache pulsation. While it is not unusual for a polygenic risk score based on migraine-associated variation to be associated with the diagnostic symptoms of migraine, the consistency of the associations and direction of effect across these symptoms suggest that migraine severity is positively correlated with higher burden of common polygenic variation. In addition to these diagnostic symptoms, we also found that migraine cases that had received treatment with triptan medication, both in the family sample and the population cohort, were associated with higher polygenic load. This again points to a higher polygenic load for individuals that are more likely to suffer from more disabling migraines since they have sought out specialist treatment.

In conclusion, our study supports the hypothesis that migraine subtypes are genetically heterogeneous diseases, and that regardless of whether they are common (i.e. MO and migraine with typical aura) or rare subtypes (i.e. FHM and SHM), common polygenic variation significantly contributes to the aggregation of the disease in families.

## URLs

GMMAT, <u>https://content.sph.harvard.edu/xlin/software.html#gmmat</u>; GWAMA, <u>http://www.well.ox.ac.uk/gwama/</u>; IMPUTE2, <u>https://mathgen.stats.ox.ac.uk/impute/imputev_2.html</u>; International Headache Genetics Consortium (IHGC), <u>http://www.headachegenetics.org/</u>; KING, <u>http://people.virginia.edu/~wc9c/KING/</u>; PLINK, <u>https://www.cog-genomics.org/plink2</u>; R, <u>https://www.r-project.org/</u>; SHAPEIT, <u>http://www.shapeit.fr</u>.

## Acknowledgements

We would like to thank the research participants and employees of 23andMe, Inc., a personal genetics company, for making this work possible.

## References

1. Agarwala, V., Flannick, J., Sunyaev, S., GoT2D Consortium & Altshuler, D. Evaluating empirical bounds on complex disease genetic architecture. Nat Genet 45, 1418–1427 (2013).

2. Khera, A. V. et al. Association of rare and common variation in the lipoprotein lipase gene with coronary artery disease. JAMA 317, 937–946 (2017).

3. Ripatti, P. et al. The contribution of GWAS loci in familial dyslipidemias. PLoS Genet 12, e1006078 (2016).

4. Stewart, W. F., Staffa, J., Lipton, R. B. & Ottman, R. Familial risk of migraine: a population-based study. Ann Neurol 41, 166–172 (1997).

5. Global Burden of Disease Study 2013 Collaborators. Global, regional, and national incidence, prevalence, and years lived with disability for 301 acute and chronic diseases and injuries in 188 countries, 1990-2013: a systematic analysis for the Global Burden of Disease Study 2013. Lancet 386, 743–800 (2015).

6. Headache Classification Committee of the International Headache Society (IHS).The International Classification of Headache Disorders, 3rd edition (beta version). Cephalalgia 33, 629–808 (2013).

7. Ophoff, R. A. et al. Familial hemiplegic migraine and episodic ataxia type-2 are caused by mutations in the Ca2+ channel gene CACNL1A4. Cell 87, 543–552 (1996).

8. De Fusco, M. et al. Haploinsufficiency of ATP1A2 encoding the Na+/K+ pump alpha2 subunit associated with familial hemiplegic migraine type 2. Nat Genet 33, 192–196 (2003).

9. Dichgans, M. et al. Mutation in the neuronal voltage-gated sodium channel SCN1A in familial hemiplegic migraine. Lancet 366, 371–377 (2005).

10. Thomsen, L. L. et al. The genetic spectrum of a population-based sample of familial hemiplegic migraine. Brain 130, 346–356 (2007).

11. Thomsen, L. L. et al. Screen for CACNA1A and ATP1A2 mutations in sporadic hemiplegic migraine patients. Cephalalgia 28, 914–921 (2008).

12. Chasman, D. I., Schürks, M. & Kurth, T. Population-based approaches to genetics of migraine. Cephalalgia 36, 692–703 (2016).

13. Anttila, V. et al. Genome-wide association study of migraine implicates a common susceptibility variant on 8q22.1. Nat Genet 42, 869–873 (2010).

14. Chasman, D. I. et al. Genome-wide association study reveals three susceptibility loci for common migraine in the general population. Nat Genet 43, 695–698 (2011).

15. Freilinger, T. et al. Genome-wide association analysis identifies susceptibility loci for migraine without aura. Nat Genet 44, 777–782 (2012).

16. Gormley, P. et al. Meta-analysis of 375,000 individuals identifies 38 susceptibility loci for migraine. Nat Genet 48, 856–866 (2016).

17. Anttila, V. et al. Genome-wide meta-analysis identifies new susceptibility loci for migraine. Nat Genet 45, 912–917 (2013).

18. Kallela, M., Wessman, M. & Färkkilä, M. Validation of a migraine-specific questionnaire for use in family studies. Eur J Neurol 8, 61–66 (2001).

19. Borodulin, K. et al. Forty-year trends in cardiovascular risk factors in Finland. Eur J Public Health 25, 539–546 (2015).

20. Lim, E. T. et al. Distribution and medical impact of loss-of-function variants in the Finnish founder population. PLoS Genet 10, e1004494 (2014).

21. Anderson, C. A. et al. Data quality control in genetic case-control association studies. Nat Protoc 5, 1564–1573 (2010).

22. Manichaikul, A. et al. Robust relationship inference in genome-wide association studies. Bioinformatics 26, 2867–2873 (2010).

23. Chheda, H. et al. Whole-genome view of the consequences of a population bottleneck using 2926 genome sequences from Finland and United Kingdom. Eur J Hum Genet 25, 477–484 (2017).

24. Surakka, I. et al. The rate of false polymorphisms introduced when imputing genotypes from global imputation panels. BioRxiv (2016). doi:10.1101/080770

25. Delaneau, O., Marchini, J. & Zagury, J.-F. A linear complexity phasing method for thousands of genomes. Nat Methods 9, 179–181 (2011).

26. Howie, B., Fuchsberger, C., Stephens, M., Marchini, J. & Abecasis, G. R. Fast and accurate genotype imputation in genome-wide association studies through pre-phasing. Nat Genet 44, 955–959 (2012).

27. Purcell, S. et al. PLINK: a tool set for whole-genome association and population-based linkage analyses. Am J Hum Genet 81, 559–575 (2007).

28. Chen, H. et al. Control for population structure and relatedness for binary traits in genetic association studies via logistic mixed models. Am J Hum Genet 98, 653–666 (2016).

29. Weiner, D. J. et al. Polygenic transmission disequilibrium confirms that common and rare variation act additively to create risk for autism spectrum disorders. Nat Genet 49, 978–985 (2017).

30. MacArthur, D. G. et al. Guidelines for investigating causality of sequence variants in human disease. Nature 508, 469–476 (2014).

31. Hiekkala, M. E. et al. The contribution of CACNA1A, ATP1A2 and SCN1A mutations in hemiplegic migraine: a clinical and genetic study in Finnish migraine families. Cephalalgia, submitted - in revision. Cephalalgia (2017).

32. Kaunisto, M. A. et al. A novel missense ATP1A2 mutation in a Finnish family with familial hemiplegic migraine type 2. Neurogenetics 5, 141–146 (2004).

33. Ferrari, M. D., Klever, R. R., Terwindt, G. M., Ayata, C. & van den Maagdenberg, A. M. J. M. Migraine pathophysiology: lessons from mouse models and human genetics. Lancet Neurol 14, 65–80 (2015).

34. Tosto, G. et al. Polygenic risk scores in familial Alzheimer disease. Neurology 88, 1180–1186 (2017).

